# Is a Simple Sensorimotor Reaction Really Simple?

**DOI:** 10.1101/2020.04.30.070706

**Authors:** Alexey A. Kulakov

## Abstract

The simple sensorimotor reaction (SSR) is widely used in psychophysiological research. It was previously shown, that the SSR latency is not constant. We studied changes in the SSR latency as a function of the waiting time from the moment of the previous response to the moment of the start of the stimulation. We performed the stimulation using light, sound and air impulse. As a response, we used a “labial sound”, a finger touch and blinking of the eyes. In all cases, where the objects of the study were humans, the SSR latency had constant and variable components. The constant SSR component was the shortest in response to closing the eyes to sound (120 ms). For “lip reaction” and finger response to sound it was 174–178 ms and 178–182 ms, respectively, but for “lip reaction” and finger response to light it was 220–226 ms. The variable SSR component represented exponential latency decay with an increase in the waiting time interval. In this case, the decay consisted of at least two components, with an apparent relaxation time in the range 30–150 ms and 600–1300 ms. Alternating stimulation of paired organs of the reception or alternating fingers reduced the apparent relaxation time of the SSR latency decay. Moreover, the latency of the human corneal reflex during eye stimulation with an air pulse also had the latency decay with three components of apparent relaxation time 9.5, 68.2 and 1,086 ms and the constant latency of 34.2 ms.

The latency of the corneal reflex in a young cat was constant and had a value of 14.6 ms. Thus, it has been shown, that the SSR latency has a complex structure and, like any conditioned reflex, is strongly influenced by the cortex. We believe that a choice is made in the centers for analysis of receiving signals from reception organs and centers sending signals to reacting organs, i.e. essentially, the SSR is also a choice reaction.

## Introduction

SSR is one of the main tools for studying many mental and psychophysiological phenomena (Jensen,1979; Niemi and Naatenen, 1981; Miller and Low,2001; Riemann and Lephart, 2002; Wood1 et al, 2015;Vidal et al,2015).Typically, the subject is given the task: in response to the appearance of a signal, for example, sound or light, press the button, or react in another way that can be registered (Niemi,1979; Jensen,1979; Miller and Low,2001).However, the subject may not find a direct relationship between the appearance of the stimulus and the response. This sometimes manifests itself in the form of a long delay in response.Meanwhile, it has recently been shown, that the time of a SSR is variable and depends on the previous reaction, or more precisely, on the time interval from the previous reaction to the onset of stimulation.The shorter the waiting interval, the longer the reaction time, and vice versa (Niemi,1979; Kulakov,2015; Kulakov,2016). It was also shown that this gradual decrease in reaction latency, depending on the waiting interval, is multicomponent and consists of 2-3 exponential decays (Kulakov,2015; Kulakov,2016;Kulakov,2018).

In this regard, it was interesting to try to find out with what processes these components of the decays are associated. This article is dedicated to this attempt.

## Methods

The studies were conducted with a man, age 68 years, as well as with a cat, age 4 months. The experiment was conducted on a hardware-computer system consisting of a computer and the “Arduino Uno” platform. This was done in order to exclude the influence of the computer on the measurement results, since the Windows operating system is multithreaded.Therefore, we placed the main program for conducting the experiment on the “Arduino Uno” platform.The program began to work from the start signal from the computer, and at the end of the experiment, the data was transferred back to the computer. The source of stimulus was the red and green LEDs in one case, inserted into dark glasses, so that when two LEDs were lit simultaneously, they would look like one. Sound stimulations were carried out via headphones, and “Arduino Uno” served as a generator. The sound frequency was 490 Hz. The sensor for receiving the signal from the subject was a microphone built into a common housing with “Arduino Uno”. In addition, we used another type of response, namely: a “lip sound”, formed under the influence of a slight air pressure during a sharp opening of the lips, a microphone was also used here.The experimental protocol for the study of latency time interval of the SSR was as follows (Fig. 1).

**Figure 1.**
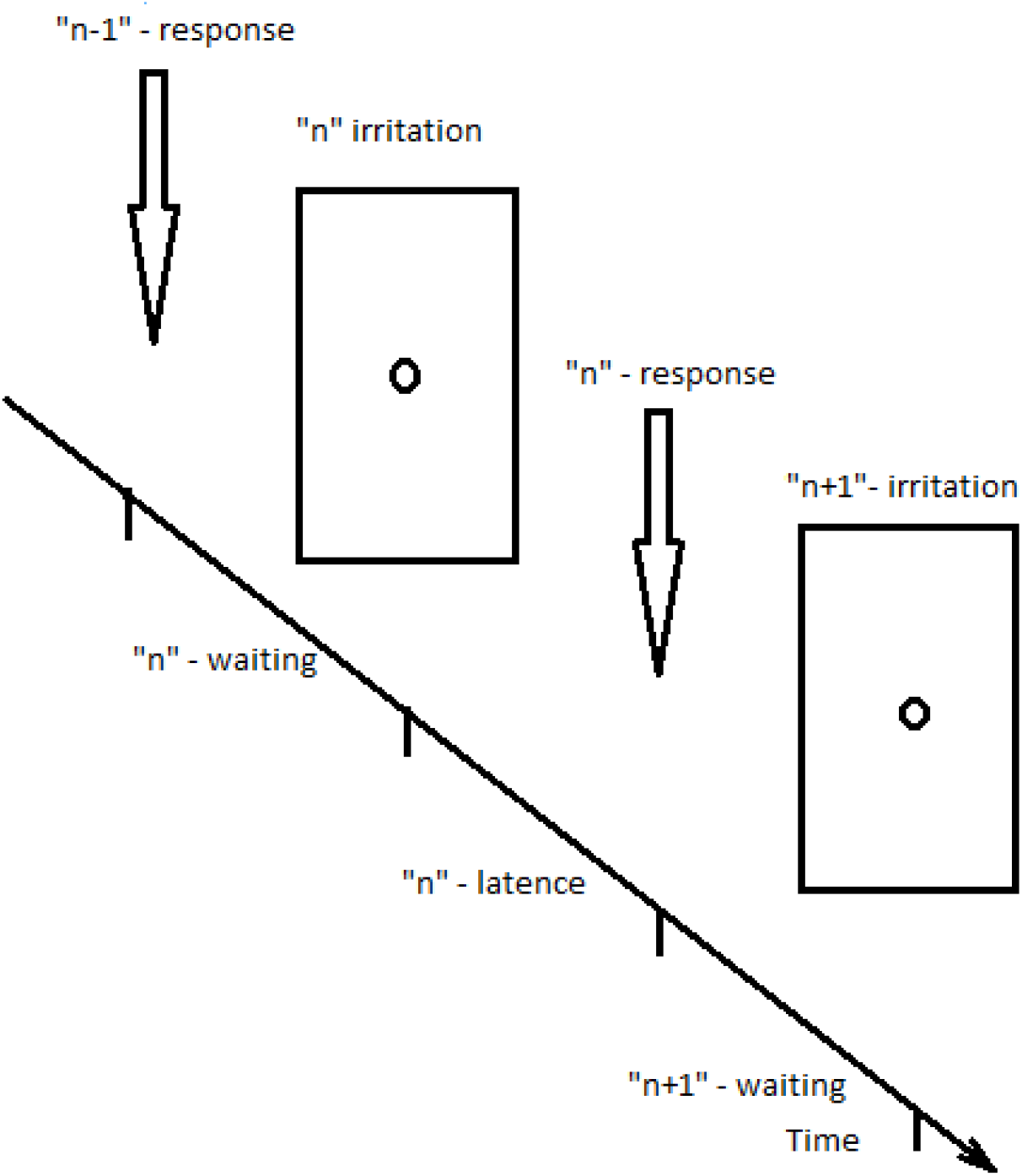
Protocol for measuring latent time of a simple sensorimotor reaction

The experiment was carried out in a cycle.Within one day, 1-2 repetitions of different options were carried out. Every day we tried to change the sequence of options in order to neutralize the phenomenon of fatigue. The total number of repetitions of one variant was 10–20.After the first presentation of the stimulus, the subject was responding with a blow of the finger on the body, or with a “lip sound”. As a result, the stimulus signal was disappearing. The reaction latency was recorded. It was followed by a predetermined waiting interval, a new stimulus signal presentation, a new response of the subject and the reaction latency recording. The experiment was repeated a predetermined number of times (usually 50) without interruption.Thus, each specified waiting time interval corresponded to its own latency interval, and in general, for the test period there was no unused time interval. It should be noted that the stimulus signal continued until the moment of response.Waiting times ranged from 3 to 5000 ms, and were grouped in the form of 5 packets of 10 signals each, and these packets could follow one after another in a given way.The result was a matrix of durations of waiting times, so that both horizontally and vertically the matrix provided almost the same average waiting time.This was done to minimize the level of influence of the time taken for a particular measurement.The total test time was approximately 80-90 seconds, depending on the subject.

The protective corneal reflex determination was carried out using a homemade installation. The solenoid pulled the steel end of the rod into itself via a solid-state relay signal from the “Arduino Uno”. The other end of the rod pushed the rubber membrane, which entered the conical cavity, and thereby pushed an air pulse through an opening with a diameter of 1.5 mm. The length of the electrical impulse from “Arduino Uno” was 20 ms. The stimulus, i.e. the air pulse, had a duration of 20-25 ms. Simultaneously with the supply of current to the solenoid, the LED was lit. The movement of the rod, LED glow, and eyelid movement was recorded with an EKEN 4K action camera with a frame rate of 120 frames per second, as well as a FireFly 8SE action camera with a frame rate of 240 frames per second. Thus, we could specify when the movement of the rod began and stopped. The selected air pulse had a velocity of the order of 2 m/s and was practically not felt at a distance of 20 cm. For a human, the direction of the air stream was carried out to the inner corner of the eye, and the test person was instructed to look outside so that the air stream would not fall on the pupil. The frame was taken as the beginning of the corneal reflex, where the second consecutive movement of the eyelid to close the eye was recorded.In the case of an experiment with a cat, she would fit on her belly and stay in that position. A pulse of air was supplied to the eye. The entire procedure lasted 45-50 seconds. In contrast to experiments on measuring the latency of SSR, time intervals between stimulation impulses were set. The size of the intervals varied from 150 to 5000 ms. As a result, the latency was determined from the moment of maximum extension of the air supply rod to the moment of second consecutive frame of movement for closing the eyelid. The interval from this last moment to the moment of subsequent maximum shift of the rod was defined as the waiting interval for subsequent measurement. As a result, the obtained waiting times varied from experience to experience, and we were forced to average both the corresponding waiting times and latencies in a series of experiments.

In addition, we determined the latent time, when the stimulus was an audio signal in the form of an impulse, and eye blinks recorded by a video camera were used as a response. The experiment was the same as for determining the corneal reflex. The results obtained were analyzed as follows.The calculated waiting times and the corresponding latencies were used to build graphs, the concept of which is given in Fig. 2. When conducting cluster analysis using the K-means method, it was shown that there are points, which are redundant for this, common for most points, dependence. These are the points mainly in the lower-left corner that have a latency of less than 100 ms, and points located on the top and right that differ in that they are far away from the main cluster of points (Fig. 2a). The first cluster indicated was characteristic of erroneous early responses.The second indicated cluster was characteristic of repeated pressures when the response was too weak and the reaction had to be repeated, or the subject delayed the reaction for unknown reasons. In conversations with the test subject, he himself noticed these errors, in the latter case he admitted that he wanted to react - but for some reason he did not succeed. Sometimes he attributed this to external, accidental stimulation.

**Figure 2.**
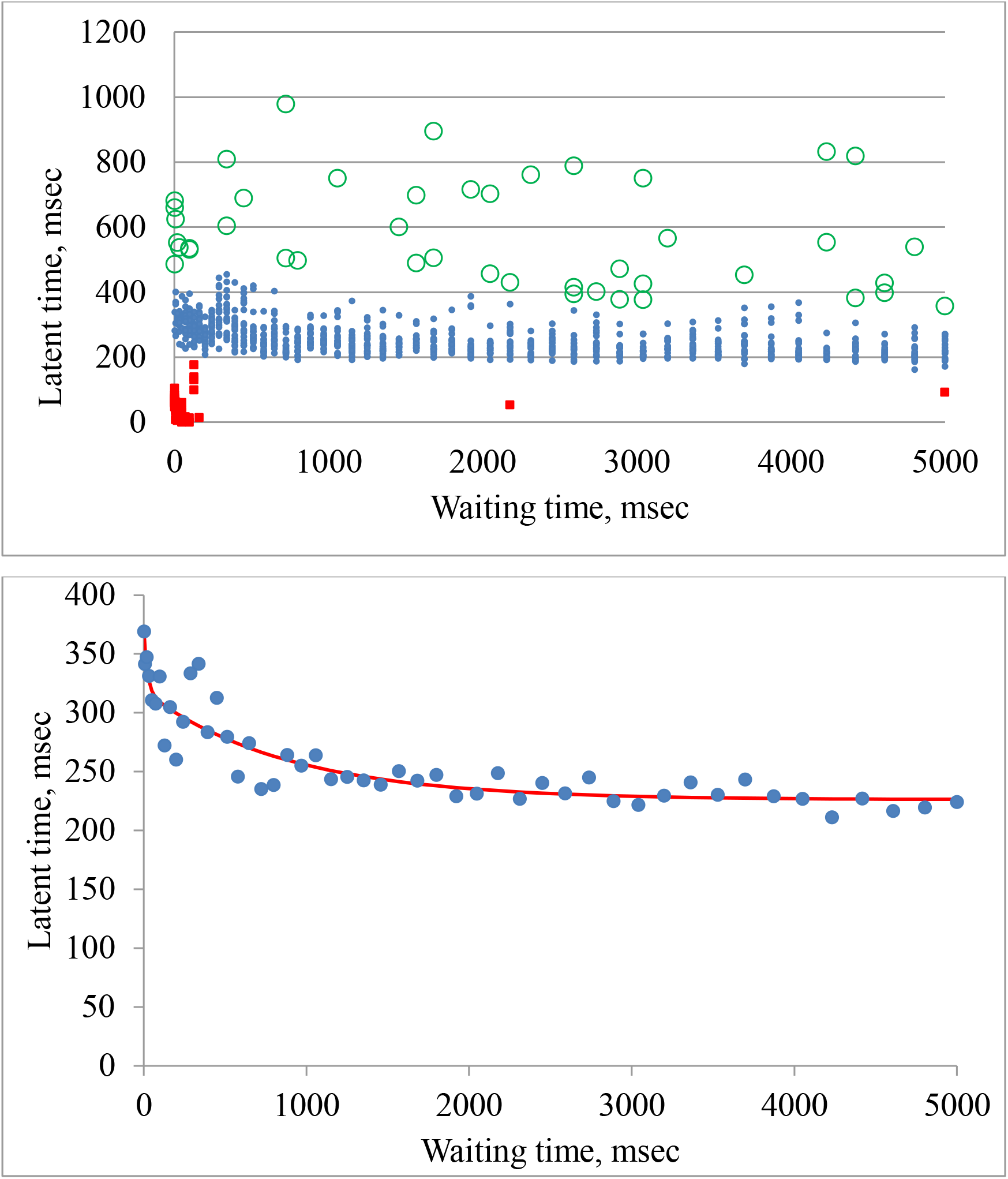
Scheme for determining the time of a simple sensorimotor reaction.

Afterwards, we averaged the latency values for the same waiting times. The resulting values are shown in Fig. 2.

A) All experimental points collected from 20 separate experiments. Large unfilled circles - discarded results due to delay, small filled squares - premature responses, small filled circles - results used for calculation. B) Circles - the average values obtained by summing the latent time values for a specific waiting time. The solid line is the result of a least squares regression.

The resulting curves then served as the basis for determining the parameters of the decay. Earlier we showed that this decay is well approximated by the sum of several exponents and the constant part [8–10], which can be expressed as the equation

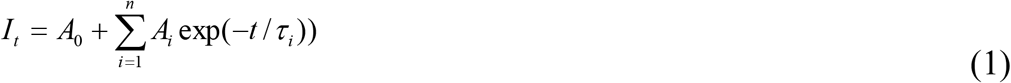

where *A_0_* is a constant latency; *А_i_* and *τ_i_* are the amplitude and relaxation time of the *i*-th component, respectively; *t* is the time interval from the previous response to stimulation; *I_t_* is the experimental value of the SSR latency; and n=4. During the calculation, the values of amplitudes and relaxation times were selected using the criterion R^2^= (*I_r_ – I_c_*)^2^, where *I_r_* and *I_c_* are the values of the ranked and calculated latencies, respectively, using the conjugate gradient method for the entire system of equations. We have set an excess number of components, since the actual number is unknown (Woods et al,2015). Components, in which the amplitudes or relaxation times were less than 10^−10^ ms, were discarded as negligible.

The parameters of the reaction of the blinking reflex were determined in a similar way, with the exception that at first the same points were averaged (relative to the interval between stimuli) both by the values of waiting times and by the latencies.This was because in these experiments the waiting time was not set, but was calculated as the time interval from the onset of stimulation to reliably visible closing of the eyelids.

## Results

We said that we were able to show that the decay in latency,occurring after the previous response,is multi-component in nature, having at least 2 components, fast and slow.It was interesting in this connection to find out whether these components can be identified with any processes.At the same time, we assumed that these components will be shorter, the shorter the reflex arc from the sensor to the motor part.In addition, we used a comparison of the alternation of perception of sensors and reacting elements, believing that the apparent relaxation time will decrease if one or the other sensor is used alternately or one or the other finger reacts. It was assumed that while the reflex arc of one of the sensors (or fingers) reacts, the reflex arc of the other sensor (or finger) is restored after the response, and the apparent relaxation time should become shorter, ideally by 2 times.The results of these trials are summarized in Tables 1 and 2. Due to the high variability, each result was obtained by processing from 12 to 22 experiments.

**Table 1.** Comparison of the parameters of a simple sensorimotor reaction with alternating or joint stimulation

| Parameters | Hip response, sound |  | Hip response, light |  |
| --- | --- | --- | --- | --- |
|  | Joint stimulation | Alternate stimulation | Joint stimulation | Alternate stimulation |
| A3 | 81,2 | 57,1 | 43,0 | 62,0 |
| T3 | 139,2 | 39,0 | 28,6 | 28,2 |
| A4 | 40,4 | 107,7 | 81, 3 | 99,9 |
| T4 | 701,9 | 288,5 | 1171,9 | 718,4 |
| A0 | 174,6 | 179,4 | 220,7 | 216,7 |
| Number of points,<br>(average error) | 1164<br>(12,5) | 768<br>(11,3) | 912<br>(12,1) | 970<br>(15) |
|  | Finger response, light stimulation |  | Finger response, sound stimulation |  |
|  | Joint stimulation | Alternate stimulation | Joint stimulation | Alternate stimulation |
| A3 | 166, 1 | 183,6 | 31,5 | 142,9 |
| T3 | 117, 2 | 99,3 | 27,0 | 6,1 |
| A4 | 3,3 | <0,00001 | 84,4 | 82,0 |
| T4 | 1736,2 | 1323,3 | 943,1 | 588,6 |
| A0 | 178,5 | 182,0 | 222,0 | 225,6 |
| Number of points,<br>(average error) | 818<br>(7,7) | 863<br>(8,8) | 874<br>(12,1) | 863<br>(12,7) |

**Table 2.** Parameters of a simple sensorimotor reaction when alternating fingers when pressed

|  | Sound, 1 finger | Sound, 2 fingers of 1 hand | Sound, 2 fingers of 2 hands<br>Light, | Light, 1 finger | Light, 2 fingers of 1 hand | Light, 2 fingers of 2 hands |
| --- | --- | --- | --- | --- | --- | --- |
| A3 | 166, 1 | 108,8 | 47,0 | 31,5 | 33,5 | 103,5 |
| T3 | 117, 2 | 95,0 | 8,6 | 27,0 | 22,5 | 3,9 |
| A4 | 3,3 | 14,8 | 90,9 | 84,4 | 79,1 | 93,9 |
| T4 | 1736,2 | 1001,9 | 380,4 | 943,1 | 1182,6 | 1000,1 |
| A0 | 178,5 | 172,7 | 181,9 | 222,0 | 239,2 | 237,0 |
| Number of points, (average error) | 818<br>(7,7) | 530 (9,1) | 491 (9,7) | 874<br>(12,1) | 525<br>(18,6) | 405<br>(13,9) |

### Analysis of Table 1

1. With simultaneous stimulation by the auditory signal of both paired sensors and reaction with one finger or lips, the constant latent time is much shorter than by stimulation with the light signal, which confirms well-known observations. In this case, responding either with the fingers or with the lips practically does not change the constant latency for the same stimulus.
2. As in our other works decays (Kulakov,2015; Kulakov,2016; Kulakov,2018), we observed a decrease in latency with an increase in the waiting time, and it turned out to be mainly a two-exponential decline.Only, in the case of a finger reacting to the alternate stimulation of both auditory analyzers, we had one component.We believe that in this case it was due to the small value of other component, which turned out to be hidden in noise.When using ranking according to the method of (Kurinov,1985) and further transformations, as in (Kulakov,2016; Kulakov,2018), this component is present.
3. In the case of simultaneous stimulation by light or sound of both paired sensors, the main components of the decay of SSR remain close and generally coincide with the data (Kulakov,2016; Kulakov,2018). The slow component has a relaxation time of about 700-1000 ms, the short one is 70-140 ms and the amplitude of both components does not exceed 180 ms.
4. When using alternating stimulation of paired analyzers, as we expected, relaxation times are reduced in comparison with simultaneous stimulation, but to a different extent.

### Analysis of Table 2

Here, our capabilities were limited by the fact that in the form of a paired response tool, we had only fingers at our disposal.However, we could vary the conditions: it was possible to react with 2 fingers on one hand, or react with 2 fingers on different hands.In this case, we used the stimulation of both pairs of analyzers: auditory or visual. As a comparison, we took corresponding experiments from Table. 1 with simultaneous stimulation and response with 1 finger.

They are also present in Table 2.You can make sure that when stimulated by light, as when pressing 2 fingers on 1 hand, or especially different hands, the fast component is greatly shortened, and the slow component is practically unchanged.At the same time, the constant component A_o_ increases somewhat. When the auditory analyzer is excited, the situation is different. When responding with the fingers of one hand, the fast component remains almost the same, while the slow component slows down. In this case, the deceleration occurs to such an extent that an additional ultrafast component of significant amplitude appears. If the fingers of different hands respond, the fast component is accelerated in the same way as in the case of responding to light, as well as the slow component.

It should be noted that a SSR is quite variable not only for different people [6], but also for an individual subject (Kulakov,2018).This makes it difficult to fully compare the data in Tables 1 and 2. Nevertheless, the results of our experiments are the basis for the assertion that a SSR is actually complex.

The next step was to use a video camera to record the response to sound. In this case, the reaction was blinking in response to an audio signal.Since we could not set the exact waiting time, we set the interval between the sound pulses (495 Hz), and the waiting time and latency were determined from the blinking results that occurred after the sound pulse, but before the next sound pulse. The results are shown in Figure 3.

**Figure 3.**
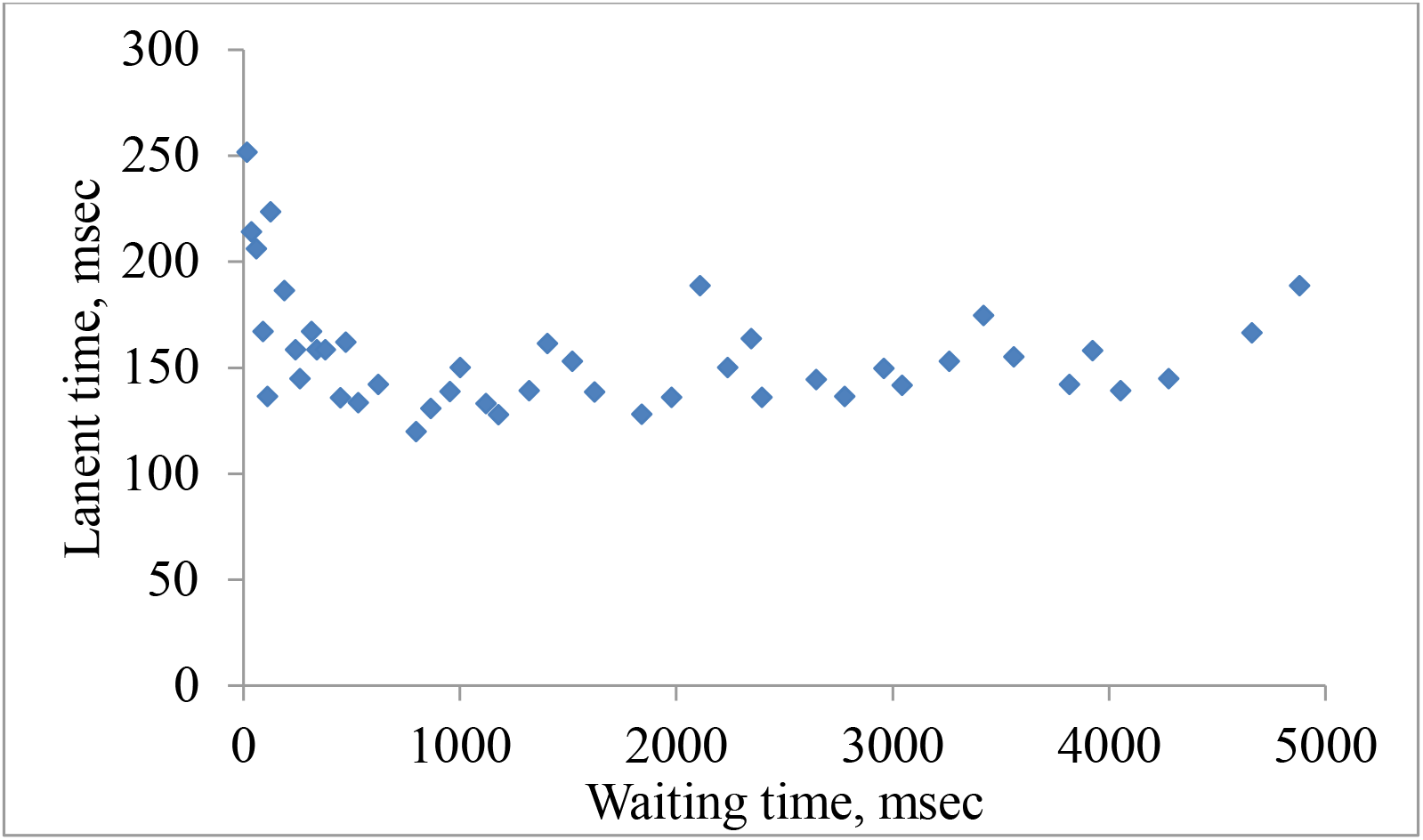
The dependence of the latent blinking time on the waiting time after a preliminary blink

As can be seen from the figure, the averaged dependence is somewhat different from the dependences for a SSR, which we observed earlier.At first, as well as for responding with lips or fingers, a decrease in latency is observed, and then, after about 1000 ms, a slow increase in latency occurs.An attempt to calculate the parameters of the latency decay using data of waiting times only up to 1000 ms gave us the constant component of the latency of 120 ms, and the relaxation constants of 99 and 814 ms, comparable to the corresponding constants in conventional response methods.

We tried to use a really SSR to study. Such is the reaction of protection (fear).This is a corneal (blinking) reflex.When the cornea is stimulated by an air pulse, involuntary closing of the eyelids occurs.This reflex is unconditional, and therefore we can expect a difference from a SSR. We studied this effect also by recording with a video camera. The results are shown in Figure 4.

**Figure 4.**
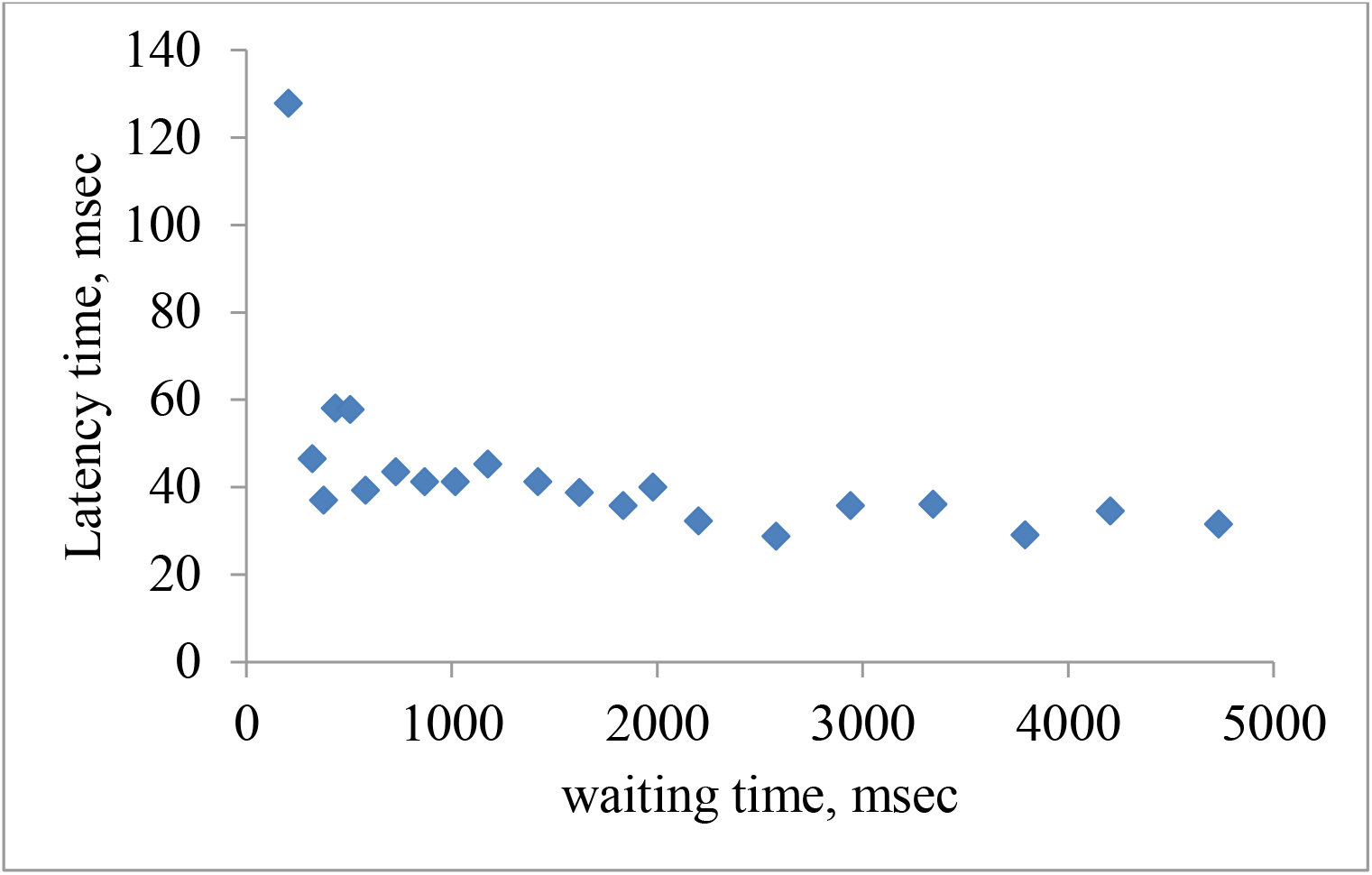
The dependence of the latent time of the corneal reflex of a person on the waiting time after the previous response

**Figure 5.**
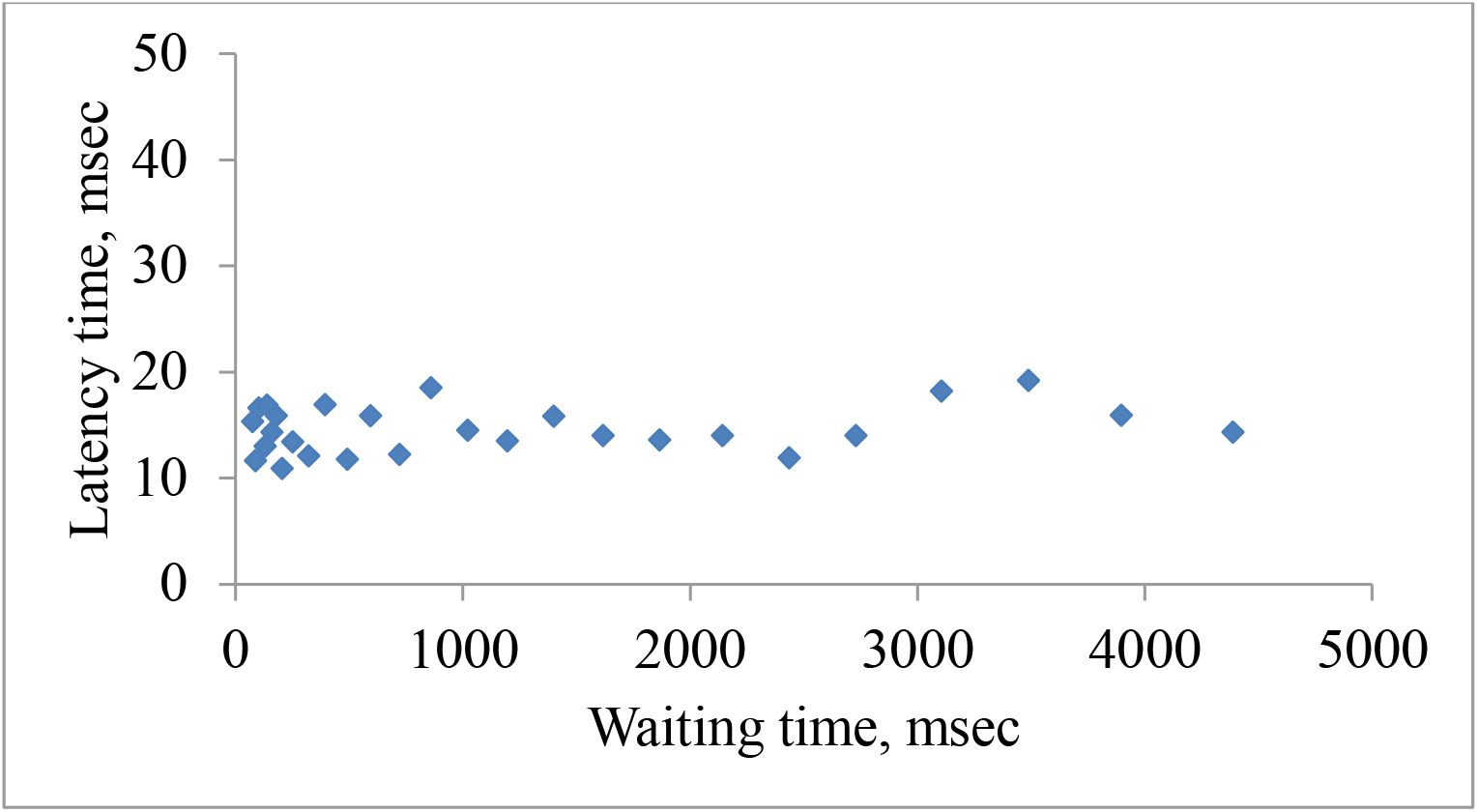
The latency of the corneal reflex of the cat as a function of the waiting time after the previous response

Analysis of the data in this figure showed that the latency of blinking after the previous blinking decreases to a different degree, similar to the latency reaction time.The decay parameters of the corneal reflex after preliminary reaction for subject K., who was our main subject, had constant latency of 34.2 ms. The slow, fast and ultrafast relaxation constants were 1086, 68.2 and 9.5 ms, respectively. Our constant latency value is consistent with the data of the authors (Medvedeva et al, 2012), who excited the blinking reflex using electrical stimulation of the supraorbital nerve.The experiments with two other subjects also showed a decrease in the latent time of the corneal effect with an increase in the waiting time.This decrease lasts more than 2000 ms, so it cannot be explained by the fact that the eye does not have time to open after stimulation.

Finally, we determined the corneal reflex on a cat, 3-months old.The Kendall coefficient in this case turned out to be +0.16, which gave reason to say that in the case of a corneal reflex for a cat, we are not dealing with a decrease in latent time after a preliminary reaction.The latency of the corneal reflex for the cat was equal to 14.6 ±4.76 ms.

## Discussion

Unfortunately, we could not find in the literature examples of studies similar to ours, with the exception of (Neemi,1979). However, the test protocol in this work was different: the waiting time there was counted from preliminary stimulation, i.e. using “foreperiod”. The vast majority of researchers use an additional signal before the main stimulation pulse, which serves either as an alarm or as a fright signal. This approach complicates understanding, because three factors have their effect on the feedback response: the time interval from the previous response to the additional signal, the time interval from the additional signal to the directly stimulating signal, and the total amount of waiting from the previous response to the directly stimulating signal.

When planning the experiments, we proceeded from several assumptions, which seem to be axioms. 1. When reducing the path of the reflex arc, the latency time should be shorter. 2. The apparent parameters of the SSR decay after a preliminary response should become shorter if alternately paired sections of the reflex arc are used, either analyzers or motor elements. This assumption is related to the fact that the currently inactive element continues to restore its original activity, which decreases during the excitation, while the other element reacts at this time.

### Consideration of the first assumption

An analysis of the experiments shows that this is partially true. Indeed, the transition from light to sound stimulation reduces a constant portion of the latency. Also, the reaction with the lips is also faster than for the fingers. If the eyelids serve as the reacting organ, then the constant part of the latent time reaches a very short time - up to 120 ms, which is almost 40 ms shorter than the response with the fingers or lips.

An even greater reduction in reaction time is observed for the corneal reflex, since the spinal component is also absent here.

At the same time, in addition to observing the corneal reflex in a cat, in all other cases there is a dependence on preliminary reaction: the longer the waiting period, the shorter the latency of a SSR, up to its constant part. Even the corneal reflex in humans also has this property. The only case – for the reaction of the eyelids to sound, we are dealing with a two-phase effect that is not clearly observable. Following the latency decay, there is again an increase in reaction latency (we did not investigate the response of the eyelids to light, because part of the time the eyes could be closed, which would distort the manifestation of the effect). However, observations of the subject showed that he had some difficulties to blink as a method of response. Next, when we asked him to perform a deliberate constant rapid blinking, it turned out that he could not stand a long, unlike, for example, a constant rapid tapping of the finger on the surface during the so-called tapping test. Roughly speaking, the eyelids were not adapted to blink quickly for a long time due to the lack of such a need. Apparently, we are dealing with fatigue here. When measuring the reaction rate for the fingers of several hundred subjects, we observed the phenomenon of fatigue in no more than 3-5%.

As we have seen, this property of responding to a stimulus is characteristic not only of a SSR, but even of the unconditional corneal reflex in humans. Only in a 4-month-old cat, this property is not observed. This allows us to say that in the reflex chain there are a number of centers, stages that affect the reflex in inhibitory manner. Thus, it can be argued that a SSR is complex, conditioned reflex, where the cortical regions of the brain are involved. This is true, because it is set verbally, and the subject may not respond for one reason or another. The author had to face such cases.

### Consideration of the second assumption

The first conclusion that we should pay attention to is the presence of inhibition when observing the SSR in all cases: when stimulating by sound or light, when reacting with the lips, fingers, and eyelids. Moreover, the nature of inhibition is almost the same: after a preliminary reaction, the latency is initially increased, and gradually decreases to a constant value, its own for each case. The second point of similarity is the presence of a slow and fast (in some cases, ultra-fast) component of latency relaxation.

Let us consider whether our results can allow us to conclude that the components of the decline in the sensorimotor reaction belong to certain structures of the nervous system.

By alternating the stimulation of paired sensors, our assumption is confirmed that the apparent relaxation constants of the decrease in the latency of the sensorimotor reaction decrease. This is evidenced by the analysis of Table 1. The degree of reduction varies depending on the conditions of the experiment. Hence, we can say that our assumption about the resulting inhibition immediately after excitation has a right to exist. Excitation from each eye goes to its center.

Therefore, when there is an alternation of irritation of one eye, then another, in the unexcited center, the ability to reactivate again is restored. Then there is a waiting period when recovery continues, i.e. the actual waiting time is really longer than in the case of simultaneous irritation of both analyzers. Therefore, the overall recovery is longer, respectively, and the apparent latency is shorter.

A similar pattern is obtained when reacting alternately with different fingers. At the same time, for the fast relaxation component, there is a clear dependence for reacting with the fingers of one hand and different hands. If the fingers of one hand react, the decrease in relaxation constants is relatively small, and for different hands is very significant. For the slow component, this dependence is for sound stimulation, but not for light stimulation, where even an increase in the slow relaxation constant occurs. In our opinion, inhibition occurs when excitation is transferred from paired centers to a single center in the case of analyzers, and from a single center to paired motor centers. We can say that in these places there is a problem of choice: to react or not to react. At the same time, there is another dichotomy, namely, in the motor centers: to transmit excitations to one hand or the other, and then another to one or another finger.

A more accurate assignment of the components, apparently, should continue. In the review (Janacsek et al,2020) for example, various assumptions about the participation of various structural components of the brain in various psychophysiological manifestations are presented: sensorimotor reaction, attention and learning. The authors (Vidal et al, 2018) suggest that motor structures and motor processes are permeable to cognitive operations; motor processes are very sensitive to the effects of cognitive operations and can contribute to the basic aspects of cognition. Comparison of various teaching methods will clarify many issues. Nevertheless, it seems to us that we were able to demonstrate the possibility of localizing certain processes responsible for the various stages of a SSR.

In conclusion, we can say that we were able to show the complexity of the SSR, and note several new approaches to its study.

## Abbreviations

SSR: simple sensorimotor reaction

## Acknowledgements

The author thanks Emil Khairullin for the translation of the article and Vadim Tkhor for editorial corrections.

## Funding

This research did not receive any specific grant from funding agencies in the public, commercial, or not-for-profit sectors.

## Notes

### Competing Interest Statement

The authors have declared no competing interest.

